# Formalin Tissue Fixation Biases Myelin-Sensitive MRI

**DOI:** 10.1101/541631

**Authors:** Alan C. Seifert, Melissa Umphlett, Marco Hefti, Mary Fowkes, Junqian Xu

**Affiliations:** Translational and Molecular Imaging Institute, Icahn School of Medicine at Mount Sinai, New York, NY USA; Department of Radiology, Icahn School of Medicine at Mount Sinai, New York, NY USA; Graduate School of Biomedical Sciences, Icahn School of Medicine at Mount Sinai, New York, NY USA; Department of Pathology, Icahn School of Medicine at Mount Sinai, New York, NY USA; Department of Neuroscience, Icahn School of Medicine at Mount Sinai, New York, NY USA

**Keywords:** Myelin, MRI, quantitative magnetization transfer (qMT), myelin water imaging (MWI), zero echo-time (ZTE) imaging, formalin fixation.

## Abstract

**Purpose:** Chemical fixatives, such as formalin, form cross-links between proteins and affect the relaxation times and diffusion properties of tissue. These fixation-induced changes likely also affect myelin density measurements produced by quantitative magnetization transfer (qMT) and myelin water imaging (MWI). In this work, we evaluate these myelin-sensitive MRI methods for fixation-induced biases.

**Methods:** We perform qMT, MWI, and D_2_O-exchanged zero echo-time (ZTE) imaging on unfixed human spinal cord tissue, and repeat these measurements after 1 day and 31 days of formalin fixation.

**Results:** The qMT bound pool fraction increased by 30.7±21.1% after 1 day of fixation and by 42.6±33.9% after 31 days of fixation. Myelin water fraction increased by 39.7±15.5% and 37.0±15.9% at these same time points, and mean T_2_ of the myelin water pool nearly doubled. Reference-normalized D_2_O-exchanged ZTE signal intensity increased by 8.17±6.03% after 31 days of fixation, but did not change significantly after 1 day of fixation. After fixation, specimen cross-sectional area decreased by approximately 5%; after correction for shrinkage, changes in D_2_O-exchanged ZTE intensity were nearly eliminated.

**Conclusion:** F and MWF are significantly increased by formalin fixation, while D_2_O-exchanged ZTE intensity is minimally affected. Changes in qMT and MWI may be due, in part, to delamination and formation of vacuoles in the myelin sheath. D_2_O-exchanged signal intensity may be altered by fixation-induced changes in myelin lipid solid-state ^1^H T_1_. We urge caution in the comparison of these measurements across subjects or specimens in different states, especially unfixed vs. fixed tissue.

## Introduction

Myelin is a biomaterial that ensheathes nerve axons, thus enhancing the fidelity and conduction speed of action potentials (1). Quantification of central nervous system myelin content has important applications to demyelinating diseases, such as characterizing demyelination and monitoring remyelination. Two widely accepted MRI methods for myelin quantification are quantitative magnetization transfer (qMT) and myelin water imaging (MWI).

Magnetization transfer (MT) exploits the exchange of magnetization between ^1^H sites on macromolecules, termed the bound pool, that cannot be directly observed by standard pulse sequences due to their broad NMR lineshape, and water ^1^H, which can be readily observed. To measure the MT effect, the longitudinal magnetization of the bound pool is first saturated by off-resonance radiofrequency (RF) irradiation, then allowed to exchange with water ^1^H, and the decrease in water ^1^H signal after this MT preparation is measured (2). Quantitative MT involves the acquisition of data at many different RF irradiation frequency offsets and amplitudes, and the fitting of these data to a signal model (3,4). This model yields the fractional size of the bound pool (F), a surrogate measure of myelin density.

Myelin water imaging discriminates between water inside the axons and in the extracellular space, and water occupying the confined spaces between layers of myelin. The latter, termed myelin water, has a shorter transverse relaxation time (T_2_) than intra/extracellular water due to its confinement by and interaction with the myelin bilayer surfaces (5,6). In MWI, the total water signal is excited and rapidly refocused by a train of 180° RF pulses, and the echoes occurring between refocusing pulses are acquired. The signal intensities in the echo-train are fitted to a sum of decaying exponential functions, with appropriate regularization, to extract the decay time constants and fractional sizes of the identifiable water pools. The portion of total signal with short T_2_ is assumed to arise from myelin water, while the portion with longer T_2_ is assumed to arise from intra/extracellular water (7). The fractional size of the short-T_2_ pool is therefore termed the myelin water fraction (MWF), another surrogate measure of myelin density.

Both qMT and MWI are relatively straightforward to implement on standard scanner hardware. Over the last decade, several *ex vivo* studies have validated their sensitivity to myelination (8–14). Consequently, both methods are widely accepted as imaging biomarkers of myelination. They are, however, indirect methods of myelin quantification, in that they interrogate the interaction of water with myelin, rather than exciting and acquiring ^1^H signal from myelin itself. This indirect nature means that care must be taken to anticipate and account for differences in the interaction of myelin and water that arise from factors other than the density of myelin. Most histologically validation studies of *ex vivo* qMT (8–10) and MWI (11–14) were performed on fixed tissue samples. Chemical fixatives, such as formalin, form cross-links between proteins (15–17), which decreases the longitudinal relaxation time (T_1_) and T_2_ of tissue water (18–21) and enhances the magnetization transfer effect (22–24). These fixation-induced changes likely bias the myelin density measurements produced by qMT and MWI (25).

Solid-state MRI methods, including zero echo-time (ZTE) (26,27) and ultra-short echo time (UTE) (28–30), are emerging techniques that have the potential to directly image and quantify the short-lived ^1^H signal arising from myelin lipids (31–34). The very presence of tissue water, however, is a significant impediment to quantification. Solid-state MRI enables the detection of myelin ^1^H signal, but does not confer specificity to this signal; water ^1^H signal is also acquired. To perform accurate quantification of myelin density *in vivo*, water ^1^H signal must be excluded from measurement. Possible methods under investigation include adiabatic inversion recovery and dual-echo subtraction (32,33,35). In contrast to indirect methods, myelin density quantification by water-suppressed solid-state MRI should be unaffected by fixation-induced changes in the interaction between myelin and water.

Another method to render solid-state MRI ^1^H signal highly specific to myelin is deuterium oxide (D_2_O)-exchange, which replaces all exchangeable ^1^H nuclei with ^2^H (deuterium, D) nuclei. ^2^H are invisible on ^1^H MRI, leaving only non-exchangeable macromolecular ^1^H nuclei to be imaged (33,36). D_2_O-exchange, however, is applicable only in *ex vivo* experiments.

In this work, we evaluated these myelin-sensitive MRI methods for fixation-induced biases. We performed qMT, MWI, and D_2_O-exchanged ZTE on unfixed human spinal cord tissue, repeated these measurements after one day and 31 days of fixation by immersion in formalin, and quantified the changes in myelin density measurements induced by tissue fixation.

## Methods

### Specimens and Equipment

Seven 20-mm segments of unfixed, unfrozen human cervical spinal cord were obtained at autopsy from four donors with no demyelinating conditions and non-neurological cause of death. The interval between time of death and time of tissue harvest was less than 48 hours.

Each 20-mm segment was divided into two 10-mm segments, each having a volume of approximately 0.7 mL. One 10-mm segment underwent qMT and MWI, and the other underwent D_2_O-exchanged ZTE imaging. The use of separate specimens is necessary to avoid decomposition by minimizing the duration of the initial unfixed imaging experiment, which must be completed before initiation of fixation (D_2_O-exchanged ZTE requires a particularly long preparation time). Specimens were stored in saline in sealed 20 mL glass scintillation vials at 4°C until time of imaging.

Imaging was performed on a 9.4 T Avance III HD micro-imaging system (Bruker, Billerica, MA) equipped with 1000 mT/m gradients and a 33-mm fixed-tune birdcage RF probe (Rapid Biomedical, Rimpar, Germany).

**Figure 1:**
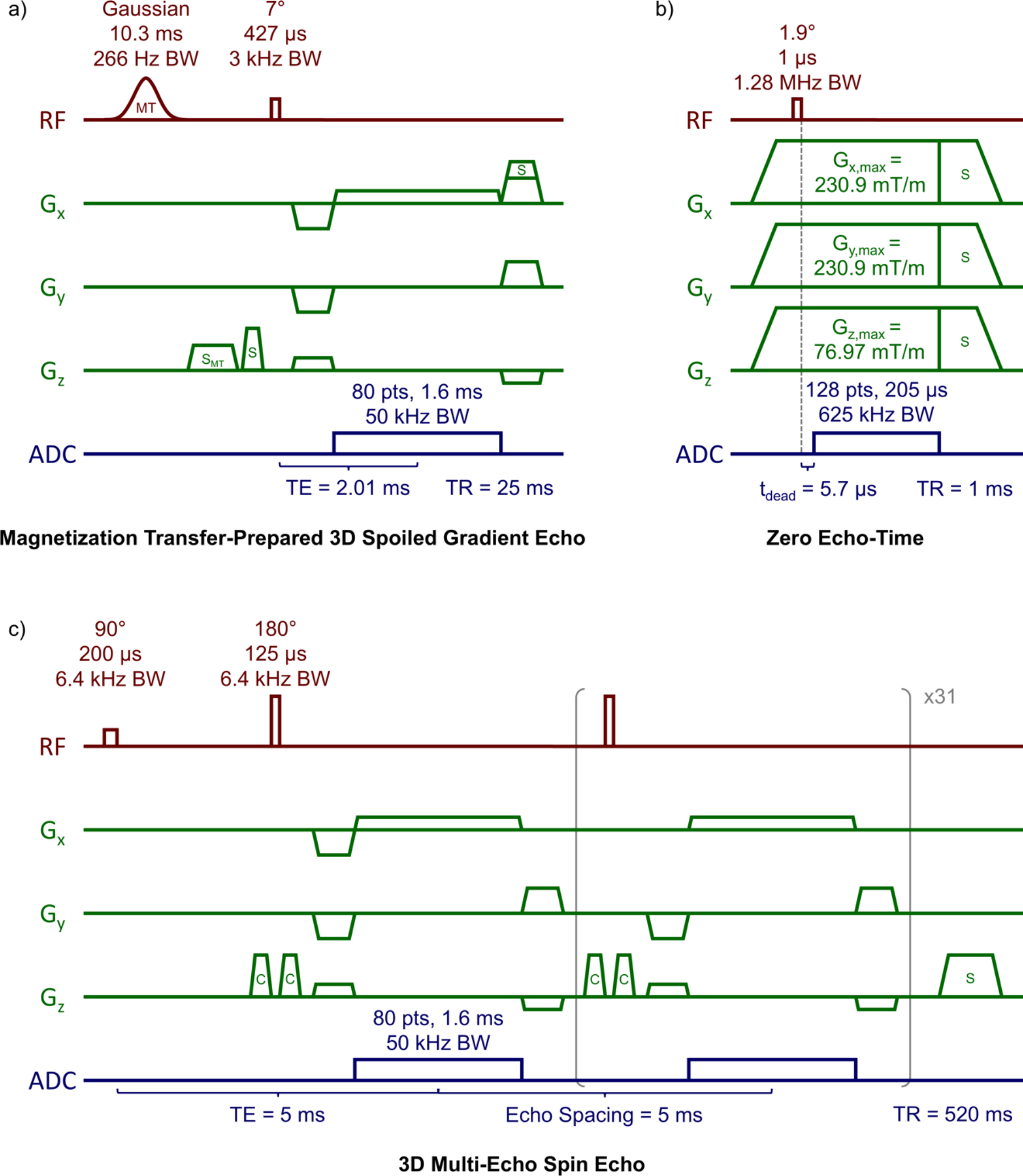
Diagrams and protocol parameters of (a) magnetization transfer (MT)-prepared 3D spoiled gradient echo, (b) zero echo-time, and (c) 3D multi-echo spin echo pulse sequences. Slice and read spoiler gradients are indicated by ‘S’, the MT-specific spoiler gradient by ‘S_MT_’, and crusher gradients are indicated by ‘C’.

Quantitative MT and MWI were performed in a single session in 20-mm glass NMR tubes at 25°C. The specimen was placed atop a 10-mm corrugated polyethylene spacer to support the 10-mm tissue specimen centrally within a 30-mm column of saline for sufficient static magnetic field (B_0_) homogeneity. B_0_ shimming was performed by at least three iterations of 3D B_0_ mapping and shim current calculation.

### Quantitative Magnetization Transfer

MT data were acquired using 61 applications of an MT-prepared 3D spoiled gradient-echo (MT-SPGR) sequence (4) with the following parameters: MT pulse shape = Gauss, MT pulse duration = 10.3 ms, MT pulse flip angles = 142° / 710° / 1388°, MT pulse bandwidth = 266.02 Hz, 20 MT pulse offsets log-spaced 234 Hz - 80 kHz, repetition time (TR) = 25 ms, excitation flip angle = 7°, excitation pulse duration = 427 μs, excitation bandwidth = 3 kHz, echo time (TE) = 2.01 ms, 2 signal averages (16 for a single MT-off acquisition), field-of-view (FOV) = 20 × 20 × 40 mm^3^, resolution = 250 × 250 × 500 μm^3^, and scan time = 6 hr 3 min.

B_0_ mapping was performed using the product field mapping protocol, utilizing a dual-echo 3D SPGR sequence with the following parameters: TR = 50 ms, TEs = 1.755 and 5.326 ms (fat and water in phase), flip angle = 20°, 4 signal averages, FOV = 20 × 20 × 40 mm^3^, resolution = 250 × 250 × 500 μm^3^, and scan time = 21 min 20 s.

Data were acquired for B_1_ mapping using six applications of a 2D SPGR sequence with the following parameters: TR = 10 s, TE = 2.54 ms, flip angle = 90°, 75°, 60°, 45°, 30°, 15°, 1 signal average each, 80 slices, slice thickness = 0.5 mm, FOV = 20 × 20 mm^2^, resolution = 250 × 250 μm^2^, and scan time = 39 min 22 s. The resulting images were fitted voxel-wise to a sine function to derive the factor by which the measured flip angle deviates from the nominal flip angle for each voxel. These B_1_ maps were then smoothed with a 3D Gaussian filter with a standard deviation equal to the voxel resolution.

Data were acquired for T_1_ relaxometry using eight applications of a 2D spoiled gradient echo sequence with the following parameters: TRs = 4920, 3360, 2280, 1560, 1080, 720, 480, and 405 ms, TE = 2.54 ms, flip angle = 90°, 1 signal average each, 80 slices at slice thickness = 0.5 mm, FOV = 20 × 20 mm^2^, resolution = 250 × 250 μm^2^, and scan time = 19 min 46 s. The resulting images were fitted voxel-wise to the steady-state spoiled gradient echo signal equation to derive the R_1_ (i.e., inverse of T_1_ relaxation time) for each voxel.

Finally, MT data were fitted voxel-wise to a two-pool MT model using qMRILab (37). Additional fitting settings and parameters were: user-provided B_1_ and B_0_ maps, R_1f_ constrained by user-provided R_1_ map, R_1r_ equal to R_1f_, model = SledPikeRP (4), lineshape = SuperLorentzian (38), and fitted parameters = F, kr, T_2f_, T_2r_. F was taken as a surrogate measure of myelin density.

### Myelin Water Imaging

Myelin water imaging data were acquired using a custom 3D multi-echo spin echo sequence (remmiRARE, Vanderbilt University Institute of Imaging Science, Nashville, TN) (39) with the following parameters: TR = 520 ms, 32 echoes, echo spacing = 5 ms, non-selective RF excitation and refocusing pulses, excitation/refocusing pulse duration = 200/125 μs, pulse bandwidths = 6.4 kHz, constant amplitude crusher gradients, 4 signal averages, FOV = 20 × 20 × 40 mm^3^, and resolution = 250 × 250 × 500 μm^3^.

To produce T_2_ spectra, echo-train signals were fitted voxel-wise by non-negative least squares to an array of 100 extended phase graph-defined exponentials under both minimum-curvature (regularization weight = 0.002) (39,40) and two-Gaussian-pool (width = 5) (6,41) constraints using the MERA toolbox (42). Additional fitting settings and parameters were: time constants spaced logarithmically from 2.5 to 500 ms, and B_1_ fitting for non-ideal refocusing pulses in the range of 150° to 210° (43,44).

Fitted peaks in the resulting T_2_ spectrum with time constants <30 ms were labeled as myelin water, and peaks with time constants >30 ms were labeled as intra/extracellular water. MWF was calculated as the integral of peaks labeled as myelin water, divided by the integral of the entire T_2_ spectrum, and MWF was taken as a surrogate measure of myelin density. All MWFs and time constants reported henceforth are generated by averaging voxel-wise values within ROIs defined on the myelin water map (45), and are taken from two-Gaussian-pool fits unless otherwise noted. Conventional T_2_ maps were also produced by voxel-wise fitting to a single mono-exponential decay.

### Deuterium-Exchanged Zero Echo-Time Imaging

Each specimen was placed in a parafilm-sealed 20-mL glass vial containing 15 mL (>20-fold volume excess) of D_2_O-saline for 24 hours at 4°C for unfixed specimens or 25°C for fixed specimens, and changed to a second sealed tube containing 15 mL of fresh D_2_O-saline for another 24 hours. This procedure results in >99.7% of H_2_O being exchanged for D_2_O (estimated as 1 − (20 + 1)^−2^, under the conservative assumption of 100% water density in the tissue). After D_2_O exchange, the specimen was placed in a parafilm-sealed 20-mL glass scintillation vial containing 15 mL of fresh D_2_O-saline, next to a 10-mm cubic piece of rubber (Hi-Polymer Eraser, Pentel, Torrance, CA), which served as a solid-state intensity reference. Localizer SPGR images (TE = 4 ms) were examined to ensure that no residual water ^1^H signal was observable in the spinal cord specimen.

Imaging was then performed at 25°C using the Bruker product ZTE pulse sequence with the following parameters: TR = 1 ms, flip angle = 2.1°, pulse duration = 1 μs, excitation BW = 1.28 MHz, transmit/receive dead time = 5.7 μs (3.56 dwells), dwell time = 1.6 μs, readout BW = 625 kHz, N = 128, readout duration = 204.8 μs, 206,742 projections (fully sampled), 16 signal averages, G_x,max_ = G_y, max_ = 231 mT/m, G_z, max_ = 77 mT/m, FOV = 64 × 64 × 192 mm^3^, voxel resolution = 250 × 250 × 750 μm^3^, and scan time = 55 min 8 s. A large FOV was required to enclose all ^1^H signal arising from the spinal cord, rubber intensity reference, and plastic RF coil structure.

ZTE image intensity, normalized to the intensity in the rubber reference sample, was taken as a quantity reflecting myelin density.

### Tissue Fixation

After pre-fixation scanning, specimens were placed in 20 mL of 10% neutral buffered formalin for 24 hours, then rinsed by serial immersion in three 20-mL volumes of saline for 24 hours each to remove residual formalin. Specimens were scanned again using the same imaging protocols. Following the 1-day fixation scans, specimens were returned to a fresh 20 mL of 10% neutral buffered formalin for an additional 30 days, rinsed, and scanned again using the same protocols as the 1-day fixation scans.

### Image Analysis

Three ROIs were drawn in a central 2-mm slab in each specimen, enclosing the dorsal column (white matter, WM), both lateral columns (WM), and both ventral horns (gray matter, GM). A fourth ROI was drawn in D_2_O-exchanged ZTE images in the rubber reference. F, MWF, and ZTE image intensity (normalized to the rubber reference intensity) values were averaged within these ROIs. To measure tissue shrinkage during the process of chemical fixation, the cross-sectional area of each spinal cord specimen was manually quantified based on the multi-echo spin echo image or the ZTE image at each time point.

### Statistical Analysis

A total of 39 confirmatory hypothesis tests were performed. Thirty-six paired two-tailed t-tests were performed to compare measurements between each pair of time points for each of the three ROIs (n = 7) and also encompassing all ROIs (n = 21). Three linear regression analyses were performed to assess the correlation of measurements across the three myelin imaging methods at all time points and in all ROIs (n = 63). The inclusion of specimen donor as a random effect decreased the model fitting quality, assessed by Akaike and Bayesian information criteria; fixed-effect linear regression was therefore used instead of mixed-effect linear regression. Statistical significance in confirmatory hypothesis tests was determined using a Bonferroni-corrected p-value threshold of 1.28 × 10^−3^ (= 0.05 / 39).

Several exploratory analyses were also performed. To correct for tissue shrinkage, myelin density measurements (F, MWF by the two-Gaussian method, and reference-normalized ZTE intensity) at each post-fixation time point were multiplied by the ratio of the respective post-fixation to pre-fixation cross-sectional area. Exploratory paired two-tailed t-tests were then repeated between each pair of time points on these shrinkage-corrected values, as well as the T_2_ relaxation times of the myelin water and intra/extracellular water pools fitted by the two-Gaussian method, MWF by the minimum-curvature method, monoexponential T_1_ and T_2_ relaxation times, and cross-sectional areas. Exploratory linear regression analyses were performed among the three shrinkage-corrected myelin density measurements, and between MWF by the two-Gaussian and minimum-curvature methods.

### Exploratory Microscopy

To support a possible explanation of the results, illustrative microscopy images were acquired on rhesus macaque and human cervical spinal cord specimens.

Luxol fast blue-stained light microscopy images of a specimen of rhesus macaque cervical spinal cord were acquired at 400x magnification. The animal was euthanized and fixed by perfusion with a solution prepared from paraformaldehyde (PFA), then the specimen was post-fixed by immersion in formalin, and stored at 4°C for over 6 months. This work was performed under a protocol approved by the institution’s IACUC.

Scanning electron microscopy images of a specimen of human cervical spinal cord were acquired at 3500x magnification. The tissue specimen was extracted at autopsy, fixed by immersion in formalin for over one year, sectioned, and post-fixed using OsO_4_. This specimen was not one of the specimens studied using MRI. Additional electron microscopy parameters were: acceleration voltage = 5000 V, emission current = 9000 nA, working distance = 23.2 mm, pixel resolution = 14.2 nm.

## Results

Representative MWI and qMT fitting results are shown in Figure 2. Differences in both MWF and F are visible as a larger area of the short-T_2_ peak in Figure 2a-b and greater magnetization transfer saturation in Figure 2c-d, respectively.

**Figure 2:**
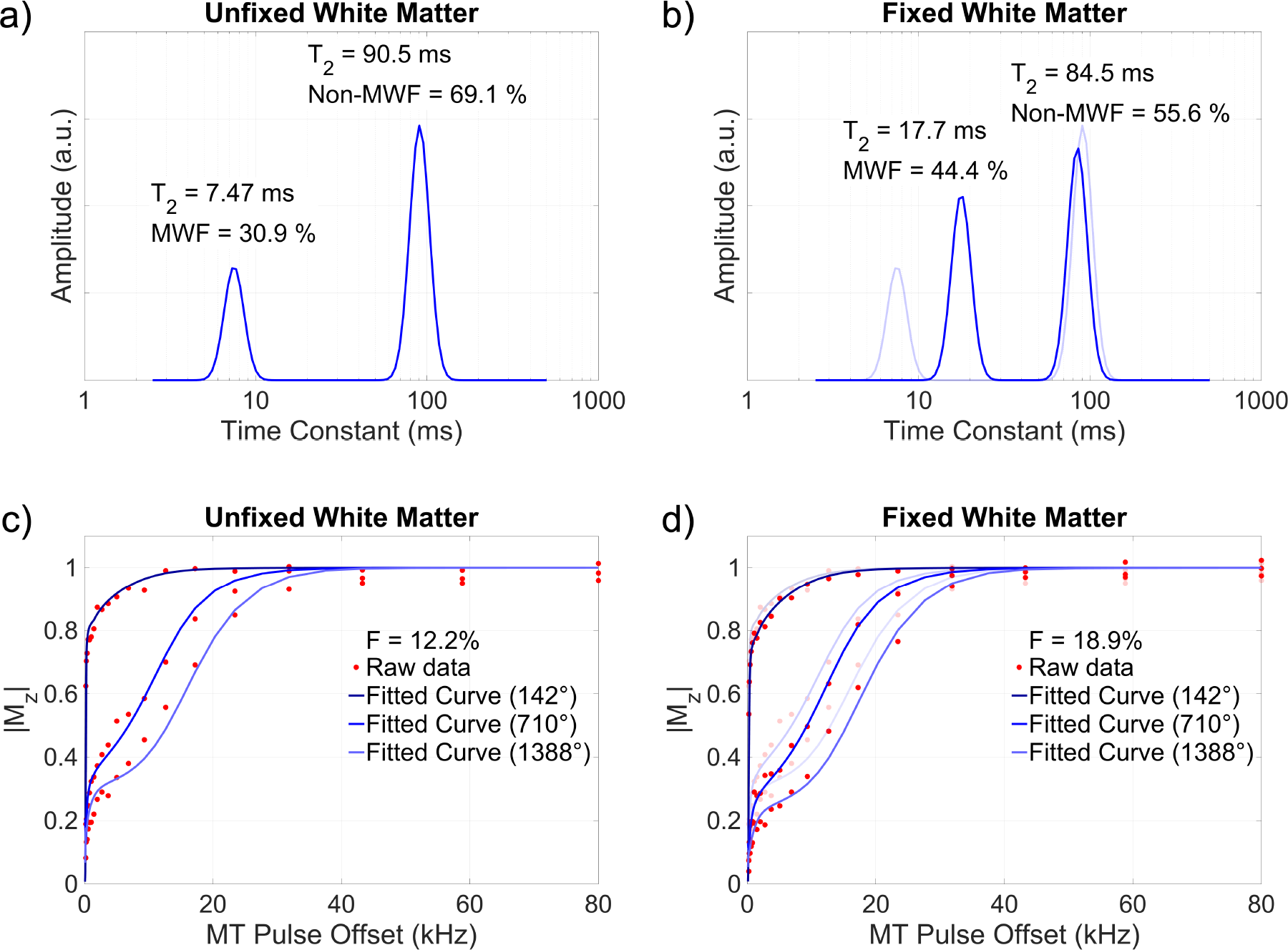
Myelin water imaging (a,b) and quantitative magnetization transfer (c,d) fitting results of a representative voxel within the dorsal white matter of an unfixed specimen (a,c) and after 31 days of fixation (b,d). After fixation, a greater fraction of total signal is identified as short-T_2_ myelin water (b vs. a), and greater off-resonance saturation yields a larger fitted quantitative magnetization transfer bound pool fraction (d vs. c). The T_2_ of the myelin water pool is also markedly increased after fixation (b). Data from panels (a) and (c) are overlaid with reduced opacity on panels (b) and (d).

Parametric maps of relaxation times and myelin density measurements in a single pair of specimens at all three time points are shown in Figure 3.

**Figure 3:**
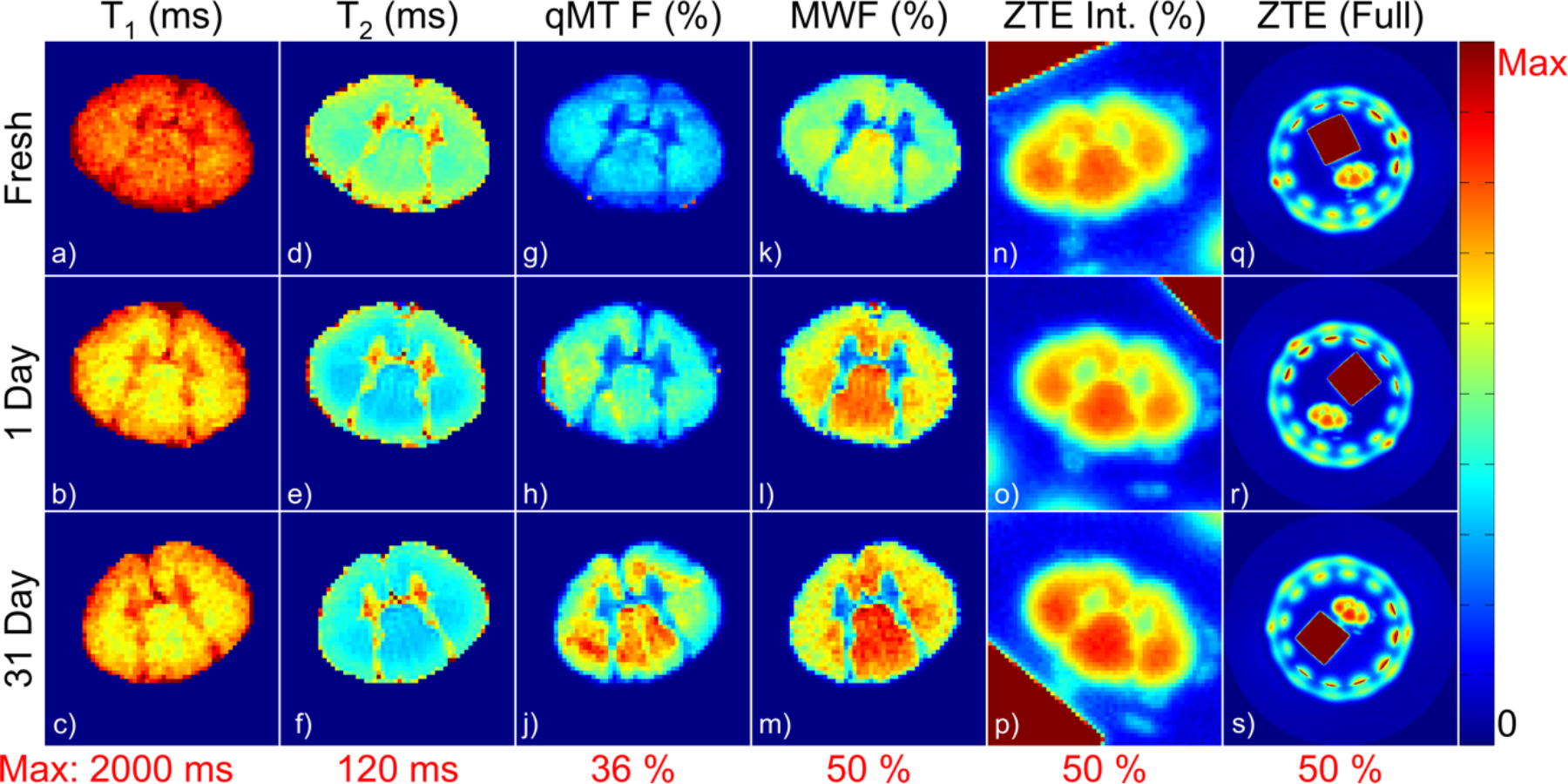
Representative parametric maps of relaxation times and myelin density measurements in an unfixed specimen and after 1 day and 31 days of fixation. All maps are displayed with the same color lookup table, scaled from 0 to the maximum value given in red below each column. The image artifact in the dorsal edge of (g) was due to Gibbs ringing in the through-slice direction arising from the plastic stage which supports the spinal cord. The ring of signal in the full-FOV ZTE images (q-s) arose from the plastic housing of the RF probe, and the high-intensity square was the rubber reference.

Changes in each of these measurements within each ROI are plotted in Figure 4a-c, and are tabulated along with summary statistics at each timepoint in Table 1. Across the three methods, F, MWF and reference-normalized D_2_O-exchanged ZTE signal intensity in the WM ROIs were consistently higher than those in the GM ROIs, as expected.

**Table 1:**
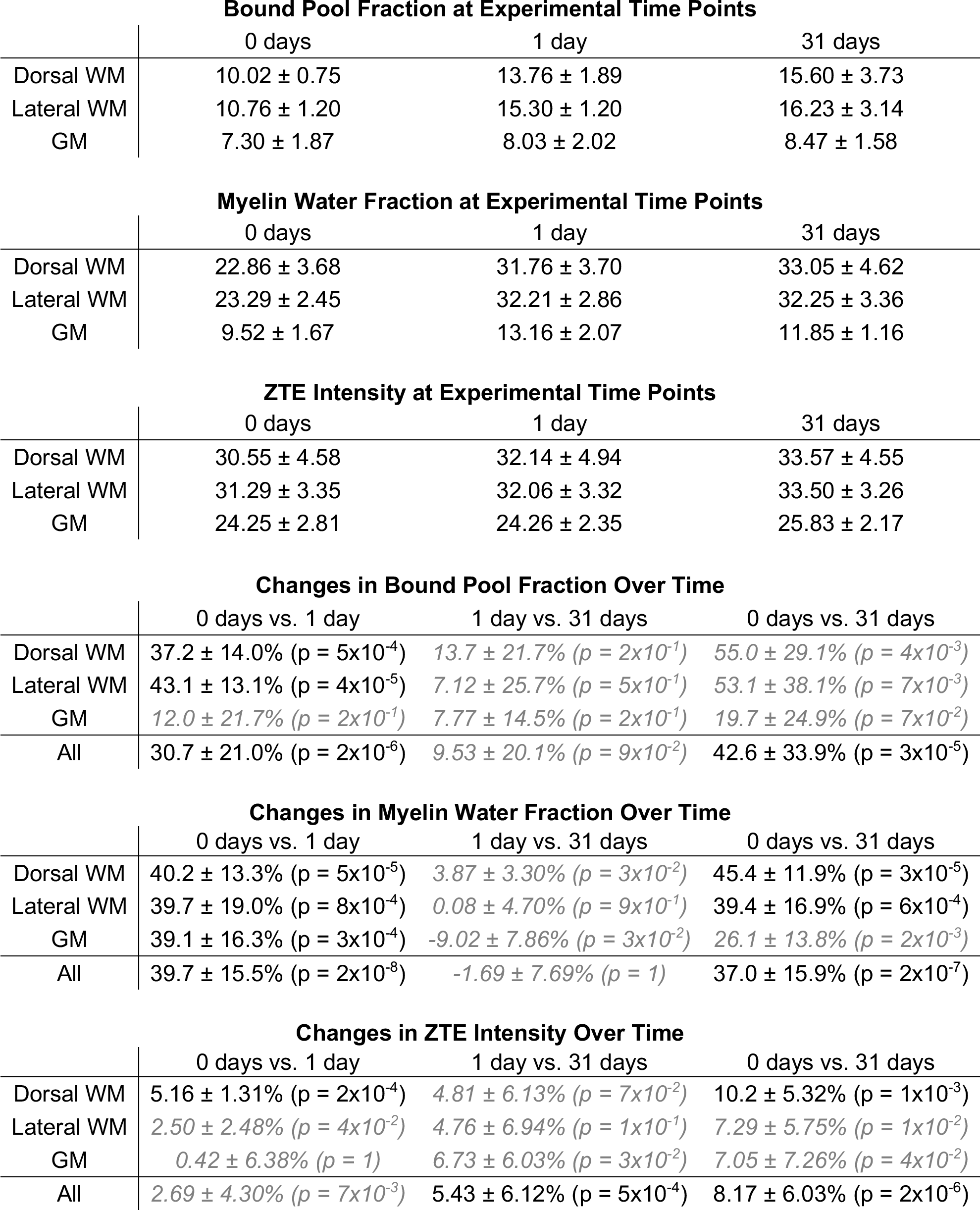
Mean ± standard deviation of myelin density measurements at each experimental time point, and changes in myelin density measurements between imaging time points expressed as percent change relative to the pre-fixation measurement. Statistical significance in paired two-tailed t-tests was determined using a Bonferroni-corrected p-value threshold of 1.28×10^−3^. Results not meeting this threshold are displayed in gray italic text. Note that comparisons in all ROIs contain n = 21 pairs of data, while comparisons in any of the three ROIs contain n = 7 pairs of data.

**Figure 4:**
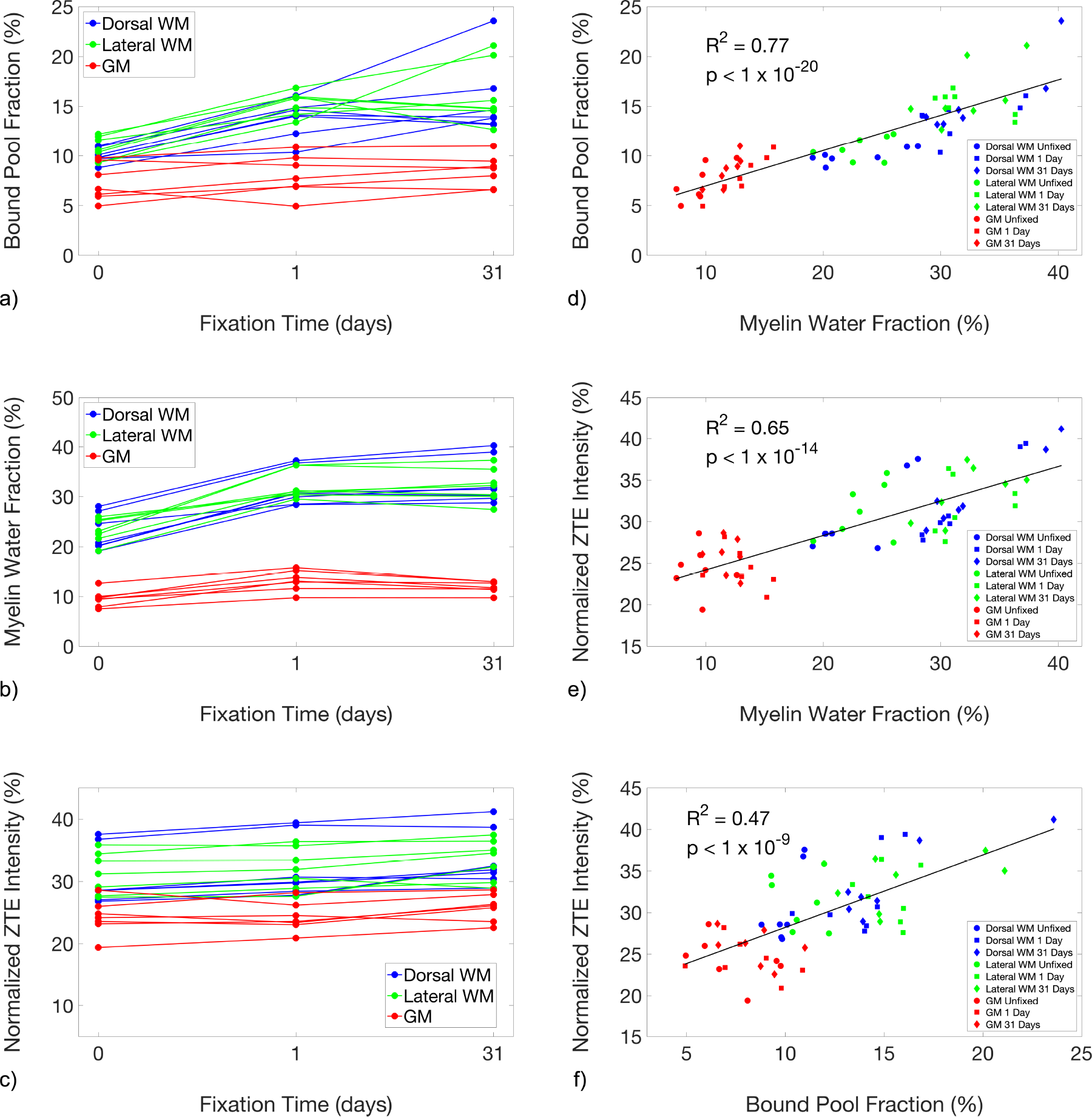
Line plots showing the changes in the quantitative magnetization transfer bound pool fraction (a), myelin water fraction (b), and reference-normalized D_2_O-exchanged ZTE signal intensity (c), of unfixed specimens and after 1 day and 31 days of fixation. Measurements produced by the three methods (d-f) were correlated. F and MWF were particularly strongly correlated (d), which suggests a common source of bias due to fixation between these two indirect measures that is not shared with D_2_O-exchanged ZTE.

Considering both WM and GM ROIs (n = 21), MWF increased rapidly in the first day of fixation, to 39.7 ± 15.5%, and remained stable in the 31-day measurement at 37.0 ± 15.9% greater than unfixed measurements. F, on the other hand, increased gradually over the full 31 days of fixation: 30.7 ± 21.0% after 1 day of fixation, and 42.6 ± 33.9% after 31 days. Reference-normalized D_2_O-exchanged ZTE signal intensities were least affected by fixation, with no significant change after 1 day of fixation and only a slight increase of 8.17 ± 6.03% (n = 21) after 31 days of fixation. Similar gradual increases were observed in both WM and GM.

In exploratory calculations, greater changes in F and MWF were observed in WM than in GM. After 1 day of fixation, F and MWF in WM ROIs (n = 14) increased by similar proportions relative to unfixed measurements: 40.1 ± 13.4%, and 39.9 ± 15.7%, respectively. After 31 days of fixation, F in WM increased further to 54.0 ± 32.6%, while MWF remained at 42.4 ± 14.4%. MWF in GM increased by 39.1 ± 16.3% after 1 day of fixation, and retreated to 26.1 ± 13.8% after 31 days. Increases in F of 12.0 ± 21.7% and 19.7 ± 24.9% were observed after 1 and 31 days of fixation, respectively.

F and MWF measurements across all ROIs and timepoints (n = 63) are strongly correlated, with R^2^ = 0.77 (p = 3×10^−21^). Correlations of F and MWF with ZTE are R^2^ = 0.47 (p = 6×10^−10^) and R^2^ = 0.65 (p = 2×10^−15^), respectively. Scatter plots comparing among the three measurements are shown in Figure 4d-f.

In exploratory analyses, the T_2_ of the myelin water pool across all WM ROIs (not including GM) in all specimens was 8.49 ± 0.97 ms in unfixed tissue, 14.5 ± 1.81 ms after 1 day of fixation (a 73.0 ± 28.5% increase), and 16.9 ± 1.01 ms after 31 days of fixation (a 100 ± 18.9% increase). The T_2_ of the intra/extracellular water pool was 87.2 ± 6.23 ms in unfixed WM, 77.0 ± 8.51 ms after 1 day of fixation (an 11.4 ± 11.2% decrease), and 81.3 ± 6.32 ms after 31 days of fixation (a 6.35 ± 9.52% decrease).

The cross-sectional areas of the spinal cords imaged by qMT and MWI (n = 7) decreased by 4.00 ± 2.94% after 1 day of fixation and 4.45 ± 8.55% after 31 days of fixation, while the cross-sectional areas of the cords imaged by D_2_O-exchanged ZTE (n = 7) decreased by 2.34 ± 1.59% after 1 day of fixation and 4.84 ± 1.25% after 31 days of fixation. After correction for this tissue shrinkage and again considering all ROIs (n = 21), F increased by 25.6 ± 20.9% after 1 day of fixation and by 37.0 ± 37.7% after 31 days of fixation; MWF increased by 33.9 ± 13.2% after 1 day of fixation and 31.2 ± 20.8% after 31 days of fixation. The change in the reference-normalized D_2_O-exchanged ZTE signal intensity nearly disappeared after correction for shrinkage: it increased by 0.30 ± 4.70% after 1 day of fixation and 2.95 ± 6.24% after 31 days of fixation. These shrinkage-corrected measurements within each ROI are plotted in Supplemental Figure 1a-c, and changes in these measurements are tabulated in Supplemental Table 3.

## Discussion

We observed substantial increases in F and MWF after fixation, and a statistically significant but small increase in D_2_O-exchanged ZTE signal intensity. The observed 54.0% increase in the F in WM from subjects without known neurodegenerative disorders is consistent with prior results. Schmierer et al. observed a 53% increase in F in normal appearing WM of brain tissue from subjects with multiple sclerosis, and 48% in multiple sclerosis lesions in the same tissues (23). Schmierer et al. also observed a 96% increase in F in the cortical GM of subjects with high cortical GM lesion load, and 49% in those with low cortical GM lesion load (24). These are considerably larger increases than the 19.7% increase we observed in spinal cord GM. Both Schmierer’s tissue samples and ours were sectioned and sufficient time was allowed for fixation, so inadequate fixative penetration was unlikely. The discrepancy may be due to differences in cellular composition (including myelinated axons in the GM) between the cortical GM sampled in Schmierer’s study and spinal cord GM.

The observed 37.0% increase (all ROIs after 31 days of fixation) in MWF is also mostly consistent with prior reports. Recent results by Chen et al. are consistent with those reported here (46), and even suggest a possible mechanism for the observed increase in MWF. In an experiment where rat spinal cords were extracted following perfusion by phosphate-buffered saline, and then fixed by immersion in formaldehyde solution, Chen et al. observed an approximately 25% increase (from ~22% to ~27%) in MWF and an increase in the T_2_ relaxation time of the myelin water pool. Most interestingly, the increase in MWF was found to be due to an increase in the absolute size of the myelin water pool. This increase in MWF was not observed in spinal cords that were fixed by perfusion with paraformaldehyde solution at the time of sacrifice; these cords yielded MWF = ~30% after perfusion fixation, and the MWF did not increase further during immersion in fixative. Both Chen’s results and our own may be explained by an expansion of the spaces between layers of the myelin sheath. These expanded spaces would contain a greater amount of water, and hence a greater fraction of the total water in the tissue. This myelin water would also have a longer T_2_, due to decreased interaction with the lipid membranes in these expanded spaces. The expansion of the myelin water pool may also contribute to the observed increase in F by facilitating greater exchange between macromolecular and water ^1^H sites.

To illustrate this hypothesized expansion of spaces, we obtained light and electron microscopic images of formalin-fixed WM (Figure 5). Delaminated myelin sheathes were observed in isolated areas in both the electron (Fig. 5d-f) and light (Fig. 5k-m) microscopic images. Cohen-Adad et al. observed this same delamination in electron microscopic images (47).

**Figure 5:**
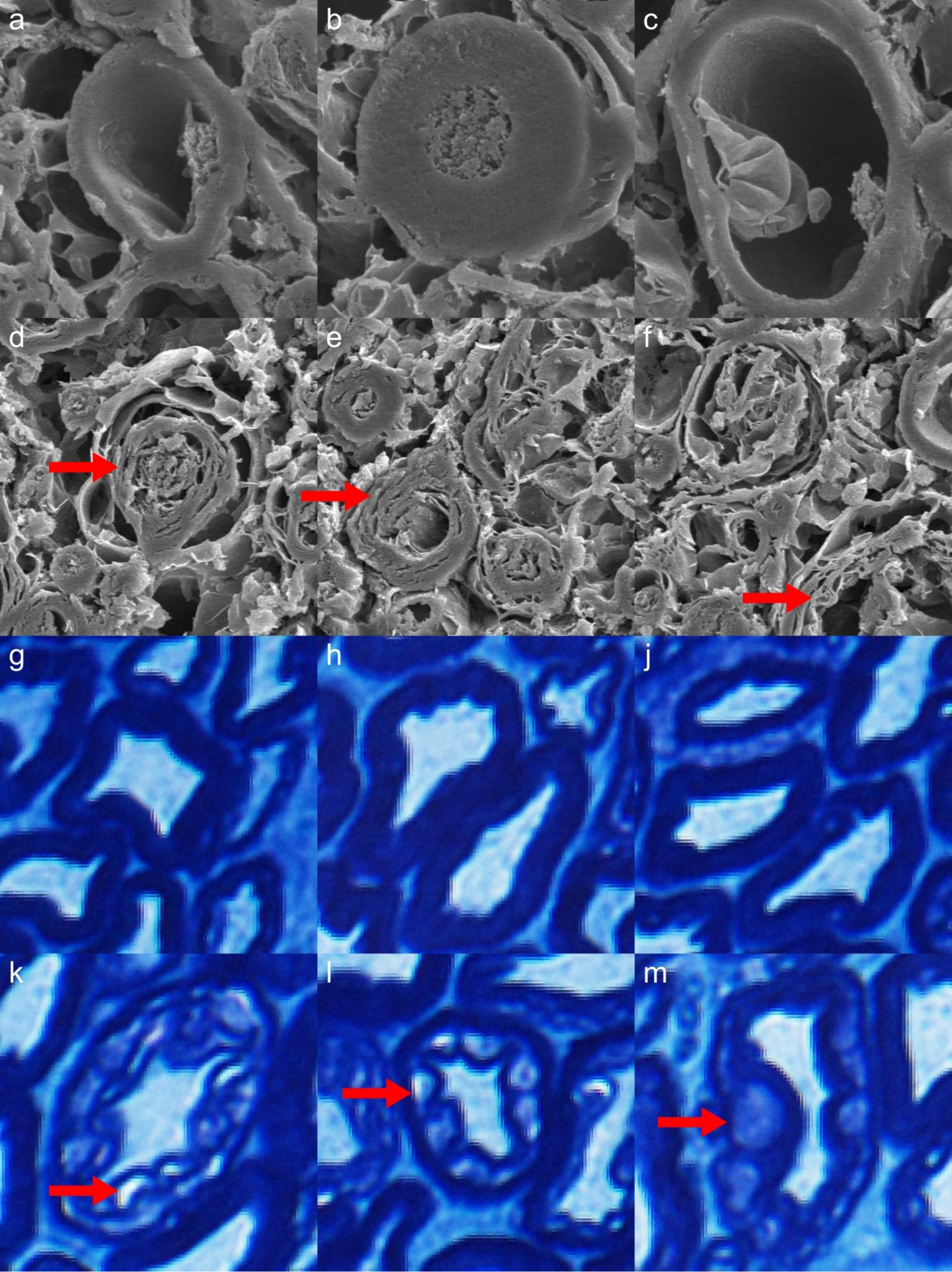
Electron microscopy (a-f) and Luxol fast blue-stained optical microscopy (g-m) images of myelinated axons from a rhesus macaque cervical spinal cord specimen fixed in formalin for 6 months. In the second (d-f) and fourth (k-m) rows, vacuoles caused by inadequate fixation by immersion in formalin are indicated by red arrows. These vacuoles may be linked to the observed increases in F and MWF in fixed tissue, relative to tightly-wrapped myelin sheathes (a-c, g-j).

Shatil et al. observed a nearly 150% increase in MWF in whole human brains fixed by immersion in formalin for approximately 40 days (25), but MWI was performed without first rinsing the fixative solution out of the tissue. The presence of formaldehyde decreases the T_1_ and T_2_ of tissue independently of its fixative effects, which may explain the much greater increase in MWF observed by Shatil et al.. Formaldehyde also diffuses slowly, needing approximately 3-5 weeks to penetrate and fix a whole brain specimen (19). The early changes in MWF observed by Shatil et al. may, therefore, be due to tissue decomposition, rather than fixation. In the present work, the small size of the spinal cord specimens ensures that fixative can rapidly penetrate the entire specimen, but the effectiveness of the fixative in preserving the structure of the myelin sheath is uncertain.

Laule et al., however, observed similar MWFs in brain tissues from subjects with multiple sclerosis before and after 2 months of fixation by immersion in formalin (see Figure 1 in (13), showing similarly-sized myelin water peaks in unfixed and fixed tissue). The heterogeneity of published MWI results suggests that the increase in MWF is not a consequence of fixation in general, but rather a consequence of fixation methods that variably or unreliably stabilize the multilamellar structure of the myelin sheath. For example, glutaraldehyde-fixed surgical nerve specimens do not generally exhibit this delamination. Glutaraldehyde, having stronger fixative properties but a lower diffusion coefficient than formaldehyde (48,49), may be able to sufficiently stabilize the myelin sheath where formaldehyde cannot. This hypothesis should be investigated further in studies pairing MWI with validation by electron microscopy.

MWI assumes that myelin water and intra/extracellular water compartments exchange magnetization slowly, compared to the range of TEs sampled, so that inter-compartmental exchange can be neglected. As inter-compartmental exchange increases, the MWF and the myelin water pool T_2_ are both expected to decrease (50–52). Inversely, if the inter-compartmental exchange rate were to decrease, the MWF and myelin water pool T2 would be expected to increase. Shepherd, et al. (53), however, found that transmembrane (intracellular-to-extracellular) water exchange rates increase by over 200% with fixation in 4% formaldehyde solution. If exchange rates between the intra/extracelluluar pools and the myelin water pool also increase with fixation, this change, considered in isolation, would cause a decrease in MWF, which is not consistent with the present results. The likely changes in intercompartmental exchange rates induced by fixation requires further investigation.

Several different methods for regularization of multi-exponential relaxation analysis have been reported in the literature and are available in the MERA software package: minimum-curvature (39,40,52) and minimum-energy (7,14,54) constraints, which impose no assumptions on number of signal pools, and a multiple-Gaussian constraint (6,41), which does impose such an assumption. We applied two of these methods to our data: minimum-curvature and multiple (two)-Gaussian. Multi-exponential T2 analysis of central nervous system WM using methods that do not constrain the number of pools generally yield only two signal components (50,55). We therefore consider a two-pool constraint to be acceptable, and we favor this method for two reasons. First, we wished to be able to consider the relaxation times of the myelin water and intra/extracellular water in combination with MWF, and this would be aided by the surety that only two pools would be fitted to each voxel’s signal. Second, after analyzing the data, we found that MWF maps produced by the two-Gaussian constraint were generally less noisy. Furthermore, MWF values averaged within ROIs drawn on voxel-wise MWF maps produced by these two regularization methods were strongly correlated: R^2^ = 0.98, p = 2×10^−51^.

The observed 8.17% increase (all ROIs after 31 days of fixation) in D_2_O-exchaged ZTE intensity after formalin fixation appears both statistically and scientifically significant at first sight. However, tissue shrinkage is a well-known effect of formalin fixation (56,57). We observed decreases in cross-sectional area of 4.64 ± 5.88%, corresponding to a reduction in volume (assuming isotropic shrinkage) of 6.88% (−15.4% to +1.9%). Due to the high degree of structural anisotropy of myelin, isotropic shrinkage was judged to be an invalid assumption, so myelin density measurements were corrected for changes in cross-sectional area. Shrinkage-corrected D_2_O-exchanged ZTE intensity was found to change by only 2.95 ± 6.24%. In a future study with a sufficiently large sample size, a statistically-significant increase in D_2_O-exchanged ZTE intensity may be found, but the increase may yet be deemed scientifically insignificant.

It is likely that the T_1_ and T_2_* relaxation times of myelin ^1^H signal are reduced by formalin fixation. Although the ZTE acquisition used a low flip angle (well below the Ernst angle) and was, therefore, proton-density weighted, a substantial decrease in T_1_ of myelin lipid ^1^H would nevertheless produce an increase in signal intensity. A decrease in T_2_* would instead result in reduced signal intensity, offsetting the effect of a decrease in T_1_, as well as increased point-spread function blurring (already significant due to the extremely short T_2_* of ~120 μs (33), much shorter than the readout duration of 204.8 μs). It is also possible that additional non-exchangeable ^1^H sites are introduced by the process of formalin fixation. The predominant source of WM solid-state ^1^H signal is methylene bridges in myelin lipids (36). Formalin fixation results in the creation of cross-links between proteins. Each cross-link, coincidentally, contains one methylene bridge, which contains two methylene ^1^H nuclei that are detectable by D_2_O-exchanged ZTE.

It is unlikely that the observed increase in D_2_O-exchaged ZTE intensity after formalin fixation is due to differences in the completeness of D_2_O exchange between fresh and fixed specimens. Fresh specimens were D_2_O-exchanged at 4°C while fixed specimens were D_2_O-exchanged at room temperature. The higher temperature would result in an approximately 80% increase in the diffusivity of water (58), which would overwhelm the approximately 27% decrease in the diffusion coefficient of water in fixed WM relative to unfixed WM (23). Incomplete exchange, therefore, would be more likely in the unfixed specimens, and, if present, would instead produce a decrease in D_2_O-exchanged ZTE intensity after fixation. Furthermore, the completeness of D_2_O exchange was confirmed in every specimen at every time point by imaging with a standard SPGR pulse sequence. In these SPGR scans, no signal was observed in either the spinal cord specimen or the D_2_O-saline solution in which the specimen and rubber intensity reference were immersed.

Our relaxometry measurements are also consistent with prior reports. We observed a 15-18% decrease in T_1_ and a 23-25% decrease in T_2_* after fixation. Dawe et al. measured a 27% decrease in T_2_* in human brain hemispheres at 3 T after fixation by immersion in formaldehyde solution (20). Raman et al. measured a ~33% decrease in T_1_ in the corpus callosum of human brain hemispheres at 3 T after 6 months of fixation by immersion in formaldehyde solution, and an approximately 15-20% decrease at one month (21).

The results presented here do not undermine the validity of F or MWF as biomarkers of myelin density in vivo or after any specific tissue preparation method. Prior work has thoroughly and convincingly proven the correspondence of these metrics to myelin concentration (8–14). In presenting these results, we instead urge caution in the comparison of measurements gained by these methods across differently-prepared specimens, especially unfixed vs. fixed specimens.

An important limitation of this study is the interval between death of the tissue donor and measurement of the pre-fixation imaging metrics. Although a strict limit of 48 hours was set between time of death and time of tissue harvest and imaging, tissue autolysis begins immediately after death, and significant changes in diffusion parameters were observed in the first three hours after death in pig brains stored at 4°C (59). Also, although D_2_O-exchanged ZTE signal intensity is minimally affected by formalin fixation, suppression of bulk tissue water by D_2_O exchange is not possible in vivo. Translation of this method to in vivo use will necessarily involve the refinement of methods to suppress tissue water by selective inversion-recovery or echo subtraction, both of which will be complicated by fixation-induced changes in the relaxation times of tissue water.

In conclusion, F and MWF are significantly increased by formalin fixation, while D_2_O-exchanged ZTE intensity is minimally affected.

## Acknowledgements

The authors gratefully acknowledge Kevin D. Harkins, Ph.D., of Vanderbilt University for providing the remmiRARE sequence, and for his assistance in modifying the protocol for ParaVision 6.0. Development of remmiRARE was supported by NIH R01-EB019980. The authors also thank Prof. Alex MacKay of the University of British Columbia for a helpful discussion on the interpretation of myelin water imaging results. This work was supported in part by NIH/NINDS K01-NS105160 and National Multiple Sclerosis Society grant PP-1705-27656. Portions of this work have previously been presented as ISMRM conference abstracts (60,61).

We are also grateful to Dr. Paula Croxson, Department of Neuroscience at Icahn School of Medicine at Mount Sinai (ISMMS) for providing the rhesus macaque spinal cord tissue; Dr. Joo-won Kim, Department of Radiology at ISMMS, for assisting with the extraction of the rhesus macaque spinal cord; Mr. William Janssen, Department of Neuroscience at ISMMS, for performing the rhesus macaque perfusion and for providing the light microscopy images; and Dr. Ronald Gordon, Department of Pathology at ISMMS, for providing the electron microscopy images.

