## Supplemental Material for "Formalin Tissue Fixation Biases Myelin-Sensitive MRI"

### Relaxometry Results

$T_1$ was 1628 ± 69 ms in unfixed white matter (n = 14), 1391 ± 83 ms after 1 day of fixation (a 14.6 ± 2.2% decrease), and 1338 ± 52 ms after 31 days of fixation (a 17.8 ± 1.8% decrease). $T_2$ was 63.35 ± 5.68 ms in unfixed white matter, 47.42 ± 4.44 ms after 1 day of fixation (a 24.8 ± 8.4% decrease), and 48.62 ± 4.34 ms after 31 days of fixation (a 22.8 ± 9.0% decrease). $T_1$ was 1734 ± 67 ms in unfixed gray matter (n = 7), 1504 ± 162 ms after 1 day of fixation (a 13.4 ± 6.9% decrease), and 1531 ± 171 ms after 31 days of fixation (an 11.9 ± 7.6% decrease). $T_2$ was 76.80 ± 14.31 ms in unfixed gray matter, 73.70 ± 12.89 ms after 1 day of fixation (a 3.48 ± 7.28% decrease), and 79.18 ± 12.30 ms after 31 days of fixation (a 3.99 ± 7.95% increase).

### Supplemental Tables and Figures

Measurements of F, MWF, and reference-normalized $D_2O$-exchanged ZTE intensity within the ROIs are tabulated before (Supplemental Table 1) and after (Supplemental Table 2) correction for tissue shrinkage. Exploratory comparisons of shrinkage-corrected measurements before and after fixation are given in Supplemental Table 3, and shrinkage-corrected measurements are plotted in Supplemental Figure 1.

**Bound Pool Fraction**

| ROI | Time (days) | Specimen 2.1 | Specimen 3.1 | Specimen 3.3 | Specimen 4.1 | Specimen 4.3 | Specimen 5.1 | Specimen 5.3 | Mean ± SD |
|---|---|---|---|---|---|---|---|---|---|
| Dor-sal WM | 0 | 9.84 | 10.97 | 10.91 | 9.72 | 8.8 | 10.09 | 9.8 | 10.02 ± 0.75 |
| | 1 | 14.01 | 16.07 | 14.86 | 10.35 | 12.26 | 14.66 | 14.11 | 13.76 ± 1.89 |
| | 31 | 13.18 | 23.58 | 16.79 | 13.84 | 14.65 | 13.22 | 13.93 | 15.60 ± 3.73 |
| Late-ral WM | 0 | 12.21 | 11.96 | 9.29 | 11.6 | 9.32 | 10.58 | 10.37 | 10.76 ± 1.20 |
| | 1 | 15.96 | 16.86 | 14.87 | 14.19 | 13.40 | 15.98 | 15.83 | 15.30 ± 1.20 |
| | 31 | 12.66 | 20.13 | 14.56 | 15.60 | 21.10 | 14.80 | 14.75 | 16.23 ± 3.14 |
| GM | 0 | 9.77 | 6.12 | 5.92 | 6.65 | 4.96 | 9.56 | 8.09 | 7.30 ± 1.87 |
| | 1 | 10.87 | 7.71 | 6.89 | 4.93 | 6.95 | 9.04 | 9.80 | 8.03 ± 2.02 |
| | 31 | 10.99 | 8.93 | 6.57 | 6.61 | 7.98 | 8.75 | 9.45 | 8.47 ± 1.58 |

**Myelin Water Fraction**

| ROI | Time (days) | Specimen 2.1 | Specimen 3.1 | Specimen 3.3 | Specimen 4.1 | Specimen 4.3 | Specimen 5.1 | Specimen 5.3 | Mean ± SD |
|---|---|---|---|---|---|---|---|---|---|
| Dor-sal WM | 0 | 24.63 | 28.06 | 27.13 | 20.75 | 20.21 | 20.15 | 19.11 | 22.86 ± 3.68 |
| | 1 | 28.49 | 37.24 | 36.76 | 29.96 | 30.78 | 30.66 | 28.41 | 31.76 ± 3.70 |
| | 31 | 29.70 | 40.26 | 38.95 | 31.89 | 31.52 | 30.24 | 28.77 | 33.05 ± 4.62 |
| Late-ral WM | 0 | 25.98 | 25.41 | 25.2 | 23.11 | 22.52 | 21.63 | 19.15 | 23.29 ± 2.49 |
| | 1 | 30.38 | 31.04 | 30.67 | 36.34 | 36.32 | 31.20 | 29.53 | 32.21 ± 2.86 |
| | 31 | 30.07 | 32.25 | 32.78 | 35.50 | 37.32 | 30.39 | 27.45 | 32.25 ± 3.36 |
| GM | 0 | 12.64 | 9.42 | 9.53 | 7.5 | 7.89 | 9.97 | 9.72 | 9.52 ± 1.67 |
| | 1 | 15.77 | 12.90 | 11.60 | 9.74 | 13.06 | 13.83 | 15.19 | 13.16 ± 2.07 |
| | 31 | 12.92 | 12.71 | 11.52 | 9.72 | 11.37 | 11.74 | 12.96 | 11.85 ± 1.16 |

**ZTE Intensity**

| ROI | Time (days) | Specimen 2.2 | Specimen 3.2 | Specimen 3.4 | Specimen 4.2 | Specimen 4.4 | Specimen 5.2 | Specimen 5.4 | Mean ± SD |
|---|---|---|---|---|---|---|---|---|---|
| Dor-sal WM | 0 | 26.81 | 37.56 | 36.76 | 28.57 | 28.55 | 28.55 | 27.03 | 30.55 ± 4.58 |
| | 1 | 27.77 | 39.41 | 39.03 | 29.89 | 29.74 | 30.69 | 28.42 | 32.14 ± 4.94 |
| | 31 | 32.47 | 41.19 | 38.68 | 31.87 | 31.41 | 30.41 | 28.93 | 33.57 ± 4.55 |
| Late-ral WM | 0 | 27.48 | 35.86 | 34.42 | 31.2 | 33.3 | 29.11 | 27.66 | 31.29 ± 3.35 |
| | 1 | 27.60 | 35.71 | 36.39 | 31.92 | 33.39 | 30.50 | 28.88 | 32.06 ± 3.32 |
| | 31 | 32.34 | 37.45 | 36.45 | 34.54 | 35.03 | 28.91 | 29.81 | 33.50 ± 3.26 |
| GM | 0 | 23.57 | 28.6 | 25.99 | 23.2 | 24.81 | 24.18 | 19.4 | 24.25 ± 2.81 |
| | 1 | 23.06 | 26.19 | 28.19 | 23.57 | 23.39 | 24.52 | 20.91 | 24.26 ± 2.35 |
| | 31 | 25.78 | 27.89 | 28.64 | 26.09 | 26.33 | 23.54 | 22.57 | 25.83 ± 2.17 |

Supplemental Table 1: Measurements of F, MWF, and reference-normalized $D_2O$-exchanged ZTE intensity without correction for tissue shrinkage. All values are in percent units.

**Bound Pool Fraction (Corrected for Tissue Shrinkage)**

| ROI | Time (day) | Specimen 2.1 | Specimen 3.1 | Specimen 3.3 | Specimen 4.1 | Specimen 4.3 | Specimen 5.1 | Specimen 5.3 | Mean ± SD |
|---|---|---|---|---|---|---|---|---|---|
| Dor-sal WM | 0 | 9.84 | 10.97 | 10.91 | 9.72 | 8.8 | 10.09 | 9.8 | 10.02 ± 0.75 |
| | 1 | 13.22 | 16.21 | 14.79 | 9.86 | 11.45 | 13.88 | 13.25 | 13.24 ± 2.09 |
| | 31 | 11.24 | 22.85 | 17.84 | 13.57 | 14.78 | 13.10 | 11.50 | 14.98 ± 4.12 |
| Late-ral WM | 0 | 12.21 | 11.96 | 9.29 | 11.6 | 9.32 | 10.58 | 10.37 | 10.76 ± 1.20 |
| | 1 | 15.06 | 17.00 | 14.80 | 13.52 | 12.52 | 15.13 | 14.87 | 14.70 ± 1.40 |
| | 31 | 10.79 | 19.50 | 15.47 | 15.29 | 21.28 | 14.66 | 12.17 | 15.60 ± 3.73 |
| GM | 0 | 9.77 | 6.12 | 5.92 | 6.65 | 4.96 | 9.56 | 8.09 | 7.30 ± 1.87 |
| | 1 | 10.26 | 7.78 | 6.86 | 4.70 | 6.49 | 8.56 | 9.21 | 7.69 ± 1.86 |
| | 31 | 9.37 | 8.65 | 6.98 | 6.48 | 8.05 | 8.67 | 7.80 | 8.00 ± 1.01 |

**Myelin Water Fraction (Corrected for Tissue Shrinkage)**

| ROI | Time (day) | Specimen 2.1 | Specimen 3.1 | Specimen 3.3 | Specimen 4.1 | Specimen 4.3 | Specimen 5.1 | Specimen 5.3 | Mean ± SD |
|---|---|---|---|---|---|---|---|---|---|
| Dor-sal WM | 0 | 24.63 | 28.06 | 27.13 | 20.75 | 20.21 | 20.15 | 19.11 | 22.86 ± 3.68 |
| | 1 | 26.88 | 37.56 | 36.58 | 28.56 | 28.75 | 29.02 | 26.69 | 30.58 ± 4.54 |
| | 31 | 25.32 | 39.01 | 41.39 | 31.26 | 31.79 | 29.96 | 23.74 | 31.78 ± 6.51 |
| Late-ral WM | 0 | 25.98 | 25.41 | 25.2 | 23.11 | 22.52 | 21.63 | 19.15 | 23.29 ± 2.45 |
| | 1 | 28.66 | 31.30 | 30.52 | 34.64 | 33.93 | 29.53 | 27.74 | 30.90 ± 2.59 |
| | 31 | 25.63 | 31.25 | 34.83 | 34.80 | 37.64 | 30.11 | 22.65 | 30.99 ± 5.36 |
| GM | 0 | 12.64 | 9.42 | 9.53 | 7.5 | 7.89 | 9.97 | 9.72 | 9.52 ± 1.67 |
| | 1 | 14.88 | 13.01 | 11.54 | 9.28 | 12.20 | 13.09 | 14.27 | 12.61 ± 1.86 |
| | 31 | 11.01 | 12.31 | 12.24 | 9.53 | 11.47 | 11.63 | 10.69 | 11.27 ± 0.97 |

**ZTE Intensity (Corrected for Tissue Shrinkage)**

| ROI | Time (day) | Specimen 2.2 | Specimen 3.2 | Specimen 3.4 | Specimen 4.2 | Specimen 4.4 | Specimen 5.2 | Specimen 5.4 | Mean ± SD |
|---|---|---|---|---|---|---|---|---|---|
| Dor-sal WM | 0 | 26.81 | 37.56 | 36.76 | 28.57 | 28.55 | 28.55 | 27.03 | 30.55 ± 4.58 |
| | 1 | 27.85 | 37.90 | 38.43 | 28.81 | 28.84 | 29.58 | 28.12 | 31.36 ± 4.68 |
| | 31 | 31.03 | 38.30 | 36.52 | 30.30 | 29.99 | 29.05 | 28.09 | 31.90 ± 3.92 |
| Late-ral WM | 0 | 27.48 | 35.86 | 34.42 | 31.2 | 33.3 | 29.11 | 27.66 | 31.29 ± 3.35 |
| | 1 | 27.68 | 34.35 | 35.83 | 30.76 | 32.38 | 29.40 | 28.57 | 31.28 ± 3.04 |
| | 31 | 30.90 | 34.83 | 34.42 | 32.83 | 33.44 | 27.62 | 28.94 | 31.86 ± 2.78 |
| GM | 0 | 23.57 | 28.6 | 25.99 | 23.2 | 24.81 | 24.18 | 19.4 | 24.25 ± 2.81 |
| | 1 | 23.13 | 25.19 | 27.76 | 22.72 | 22.68 | 23.63 | 20.69 | 23.69 ± 2.24 |
| | 31 | 24.63 | 25.94 | 27.04 | 24.80 | 25.14 | 22.49 | 21.91 | 24.56 ± 1.82 |

Supplemental Table 2: Measurements of F, MWF, and reference-normalized $D_2O$-exchanged ZTE intensity after correction for tissue shrinkage. All values are in percent units.

**Bound Pool Fraction (Corrected for Tissue Shrinkage)**

| | 0 days vs. 1 day | 1 day vs. 31 days | 0 days vs. 31 days |
|---|---|---|---|
| Dorsal WM | *31.7 ± 14.4% (p = 2x10$^{-3}$)* | *13.5 ± 24.2% (p = 2x10$^{-1}$)* | *48.7 ± 33.6% (p = 1x10$^{-2}$)* |
| Lateral WM | 37.4 ± 14.2% (p = 2x10$^{-4}$) | *7.54 ± 31.8% (p = 6x10$^{-1}$)* | *47.7 ± 44.5% (p = 2x10$^{-2}$)* |
| GM | *7.53 ± 21.3% (p = 5x10$^{-1}$)* | *7.48 ± 18.5% (p = 5x10$^{-1}$)* | *14.6 ± 27.5% (p = 3x10$^{-1}$)* |
| All | 25.6 ± 20.9% (p = 2x10$^{-5}$) | *9.49 ± 14.3% (p = 2x10$^{-1}$)* | 37.0 ± 37.7% (p = 4x10$^{-4}$) |

**Myelin Water Fraction (Corrected for Tissue Shrinkage)**

| | 0 days vs. 1 day | 1 day vs. 31 days | 0 days vs. 31 days |
|---|---|---|---|
| Dorsal WM | 34.5 ± 11.8% (p = 2x10$^{-4}$) | *3.35 ± 8.91% (p = 3x10$^{-1}$)* | *39.3 ± 19.5% (p = 2x10$^{-3}$)* |
| Lateral WM | 33.8 ± 15.8% (p = 8x10$^{-4}$) | *-0.23 ± 11.3% (p = 9x10$^{-1}$)* | *33.6 ± 22.5% (p = 8x10$^{-3}$)* |
| GM | 33.4 ± 13.8% (p = 3x10$^{-4}$) | *-9.26 ± 12.5% (p = 9x10$^{-2}$)* | *20.8 ± 18.6% (p = 4x10$^{-2}$)* |
| All | 33.9 ± 13.2% (p = 3x10$^{-8}$) | *-2.05 ± 11.8% (p = 1)* | 31.2 ± 20.8% (p = 2x10$^{-5}$) |

**ZTE Intensity (Corrected for Tissue Shrinkage)**

| | 0 days vs. 1 day | 1 day vs. 31 days | 0 days vs. 31 days |
|---|---|---|---|
| Dorsal WM | *2.69 ± 1.68% (p = 7x10$^{-3}$)* | *2.10 ± 5.33% (p = 4x10$^{-1}$)* | *4.83 ± 5.30% (p = 5x10$^{-2}$)* |
| Lateral WM | *0.11 ± 3.07% (p = 1)* | *2.05 ± 6.03% (p = 4x10$^{-1}$)* | *2.11 ± 5.89% (p = 4x10$^{-1}$)* |
| GM | *-1.90 ± 7.00% (p = 4x10$^{-1}$)* | *4.00 ± 5.86% (p = 1x10$^{-1}$)* | *1.92 ± 7.79% (p = 7x10$^{-1}$)* |
| All | *0.30 ± 4.70% (p = 8x10$^{-1}$)* | *2.72 ± 5.53% (p = 6x10$^{-2}$)* | *2.95 ± 6.24% (p = 5x10$^{-2}$)* |

Supplemental Table 3: Changes in myelin density measurements between imaging time points, after correction for tissue shrinkage, expressed as percent change relative to the pre-fixation measurement, with p-values. Although these statistical tests are exploratory, not confirmatory, changes with p-values greater than 1.28x10$^{-3}$ are displayed in gray italic text.

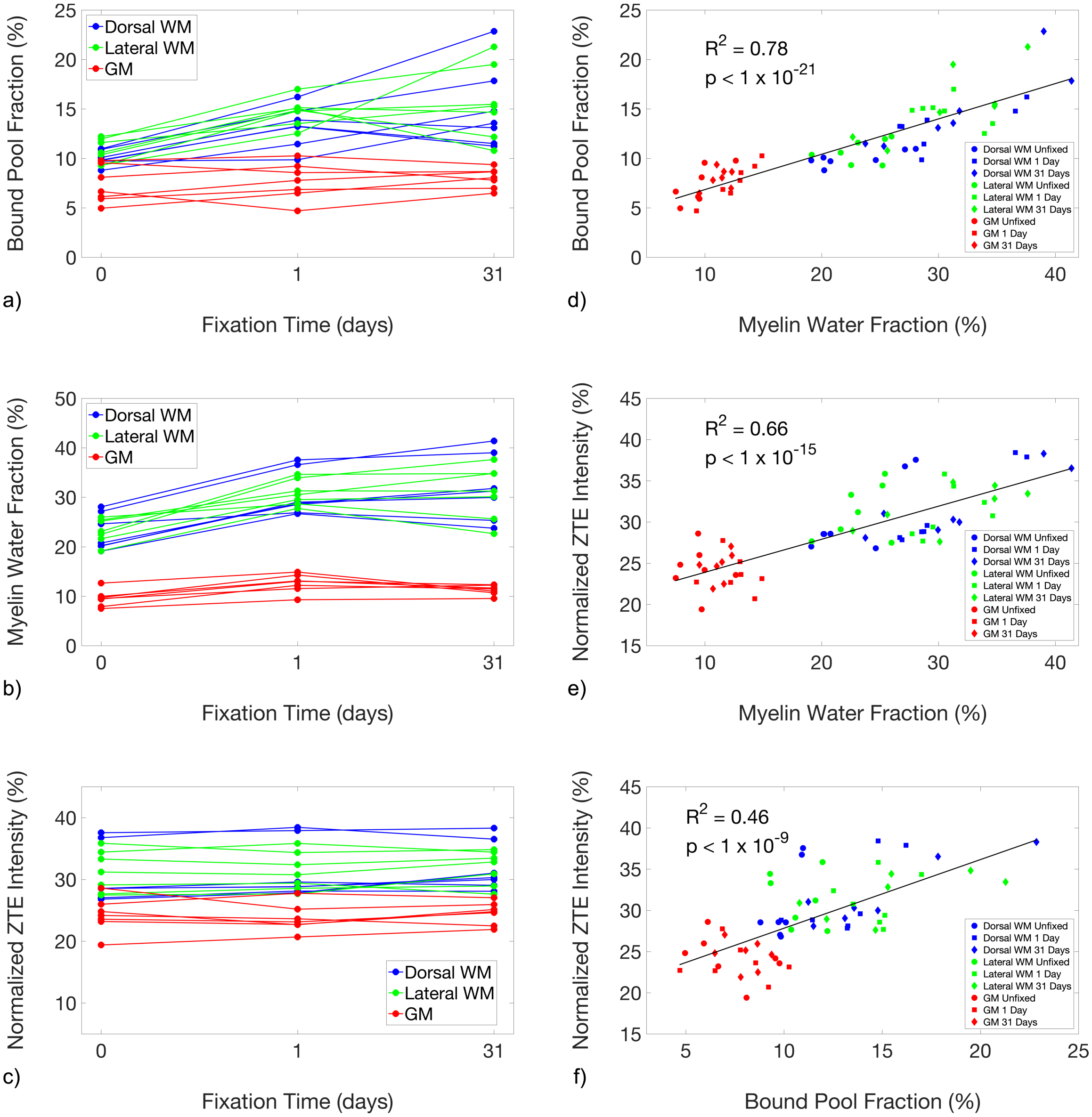


Supplemental Figure 1: Line plots showing the changes in the quantitative magnetization transfer bound pool fraction (a), myelin water fraction (b), and reference-normalized $D_2O$-exchanged ZTE signal intensity (c), of unfixed specimens and after 1 day and 31 days of fixation, after correcting for tissue shrinkage due to fixation. Shrinkage correction removes a large amount of the observed increase in $D_2O$-exchanged ZTE intensity, but the inter-measurement correlations (d-f) are not significantly altered.
